# *IN VITRO* 5-LOX INHIBITORY POTENTIAL AND ANTIOXIDANT ACTIVITY OF NEW ISOXAZOLE DERIVATIVES

**DOI:** 10.1101/2024.01.08.574589

**Authors:** Waqas Alam, Haroon Khan, Muhammad Saeed Jan, Maria Daglia

## Abstract

5-Lipoxygenase (5-LOX) is a key enzyme involved in the biosynthesis of pro-inflammatory leukotrienes, leading to asthma. Developing potent 5-LOX inhibitors are highly attractive. In this research the previously synthesized isoxazole derivatives has been investigated against 5-Lox inhibitory and antioxidant in vitro assay. The percent inhibition for **C3** were 89.93±1.73, 85.94±0.91, 81.90±1.32, 77.51±0.59 and 74.80±1.41 at various concentrations with IC_50_ of 8.47 µM. The investigated compounds **C5** also exhibited good 5-LOX inhibitory effect. The non-significant percent inhibition for **C5** were demonstrated as 91.30±1.42, 87.78±0.45, 84.44±0.86, 79.72±1.89 and 75.29±1.64 at various concentration (1000-62.5 µg/ml). The IC_50_ demonstrated for **C5** was 10.48. Among the 10 synthesized compounds, the potential 5-LOX inhibitory effect was reported for **C6**. The most potent compound which showed excellent free radical scavenging effect was **C3** with different percent inhibitions of 92.51±0.62, 87.65±0.70, 82.25±0.55, 79.37±0.69 and 75.72±0.51 having IC_50_ value of 10.96 µM. The next most potent antioxidant activity was reported for **C5** which non-significantly showed free radical scavenging effect. The DPPH percent inhibition reported for **C5** was 92.63±0.64, 88.45±0.55, 83.53±0.41, 79.42±0.46 and 76.10±0.64 at different ranges of (1000, 500, 250, 125 and 62.5 µg/mL) concentrations, respectively. The IC_50_ value observed for **C5** was 13.12 µM. Compound **C6** also showed potent dose dependent antioxidant effect with IC_50_ value of 18.87 µM having percent inhibition of 91.63±0.55, 88.45±0.49, 83.53±0.45, 78.42±0.66 and 73.72±0.64 at concentration 1000-62.5 µg/mL respectively.

Among the tested compounds, **C6** was found most potent which showed significant 5-LOX percent inhibition assay and also reported the minimum IC_50_ value comparable to the reference drug. The *in vitro* 5-Lox enzymes inhibition assays of **C5** and **C3** also showed non-significant percent inhibition and good potency next to **C6**. We concluded that amongst the investigated designed molecules the **C3** was found best potent and showed significant dose dependent antioxidant activity against DPPH screening. The IC_50_ value reported for **C3** was found good as compared to standard drug. Moreover, **C5** and **C6** also showed excellent free radical scavenging effect against DPPH assay.

## 1 INTRODUCTION

Lipoxygenase (LOX) enzymes are discovered in many plants and animals, but not in bacteria or yeast. The enzyme 5-lipoxygenase (5-LOX) is responsible for the biosynthesis of leukotrienes from the precursor AA (Biringer, 2020). Fatty acids are oxidized by non-heme iron enzymes like LOX to produce lipid hydroperoxides (Lenis et al., 2010). The presence of 5-LOX and its interaction with arachidonic acid is a vital first phase in the production of leukotrienes (LTs). In typical cellular structures, the release of arachidonic acid (AA) is primarily facilitated by endogenous phospholipids. This process is thought to be mediated by cytosolic phospholipases, which work in conjunction with many other enzymes, notably LT-converting phospholipase A2. Lipid peroxides are mostly formed as a result of LOXs (Gaschler and Stockwell, 2017).

The pathophysiological role of 5-LOX in respiratory and cardiovascular diseases has been revealed by subsequent studies that have examined the enzymatic pathway of 5-LOX. 5-LOX is a non-heme metalloenzyme that has iron at its active site; iron is present in ferrous form during rest and in ferric form during active state; hydroperoxidation of lipids due to physiological stress may activate 5-LOX; it has been demonstrated that the activated enzyme in the cytoplasm is translocated to the nuclear membrane upon Ca2+ binding and/or by phosphorylation (Muthuraman et al., 2019).

Leukotrienes are known to be lipid mediators for inflammatory response with major roles in cardiovascular diseases (Hofmann and Steinhilber, 2013). Results from recent studies also suggest the role of leukotrienes in prostate cancer, osteoporosis and in certain forms of leukemia. 5-LOX is validated as a potential drug target for treating asthma, rheumatoid arthritis, osteoarthritis, allergic, cardiovascular diseases and certain types of cancer. For more than three decades considerable progress have been made towards finding more potent 5-LOX inhibitors. Several natural products, their analogs and new scaffolds have been identified as potent 5-LOX inhibitors, to name a few, nordihydroguaretic acid (a polyphenolic compound isolated from creosote plant), curcumin (a polyphenolic compound found in turmeric), eugenol (a polyphenolic compound found in clove) and quercetin (a flavone found in garlic) (Prasad et al., 2004), Hyperforin (a phytochemical isolated from *Hypericum perforatum*) (Feißt et al., 2009), zileuton, ICI207968, ICID2138 (McMillan and Walker, 1992), 3-tridecyl-4,5-dimethoxybenzene-1,2-diol and Licofelone (Scheme 1). Based on the inhibitory mode of action, the 5-LOX inhibitors are categorized into four types (Fischer et al., 2007), (i) redox inhibitors (redox-active compounds interrupts the redox cycle of the enzyme), (ii) iron-ligand inhibitors with weak redox active properties (inhibit by chelating to active site iron), (iii) non-redox inhibitors (compete with the substrate, arachidonic acid at the active site), and (iv) allosteric inhibitors (inhibition by binding other than active site). Despite considerable efforts made towards the development of efficient drugs that target 5-LOX enzyme, zileuton remains the only clinically approved 5-LOX inhibitor in the treatment of asthma but its use is limited due to severe side effects such as, weak potency, liver toxicity, and an unfavorable pharmacokinetic profile with a short half-life (Sinha et al., 2016, Tomy et al., 2015). The antioxidant potential of organic compounds must be evaluated as they are used in food, medicine, and cosmetics (Halliwell, 1996). Reactive species are produced in living systems by a variety of metabolic processes and stressful circumstances. They are primarily reactive oxygen species (ROS). Raised levels of ROS may alter biomolecules’ activities and impair their structural integrity, which can cause cellular malfunction and even premature death of cells. An rise in ROS over time can lead to oxidative stress at the systemic level, which manifests as a number of health issues including cancer, inflammation, age-related disorders, and heart problems (Grune et al., 2001, Noguchi and Niki, 2000). The most common and simple colorimetric technique for assessing the antioxidant capabilities of pure molecules is the DPPH assay, which is frequently used to determine how well a certain antioxidant molecule scavenges free radicals (Cheng et al., 2006). Researchers have reported antioxidant potential of isoxazole derivatives. Bhatia et al., has reported the antioxidant effect of Indole-Functionalized Isoxazoles derivatives. They found that in the DPPH test, the created compounds showed variable capacity to scavenge free radicals. The isoxazole ring of this molecule is considered as potent antioxidant moiety. The compounds’ capability to inhibit free radicals was significantly influenced by the substituted sequence at the phenyl ring connected to the isoxazole group (Bhatia et al., 2022). Therefore, there is a strong need for the development of safer and more potent 5-LOX inhibitors. In the present study we have investigated the already synthesized ten pyrimidine isoxazole derivatives and evaluated their 5-LOX inhibition and antioxidant *in vitro* activities.

## 2 MATERIALS AND METHODS

### 2.1 Chemicals

Leukotriene (CAS 71160-24-2, Icatibant (CAS 138614-30-9). DPPH, Montelukast (Montiget by Getz pharma).

### 2.2 Instruments

UV-visible spectrophotometer Model-300BB, UV spectrophotometer Model UV-1800, Japan, Microplate reader-Type ABS 220-4, Biotek, Electronic balance Model-ABS 220-4, kern &Sohn GmbH, Incubator-Model 100-800, Digital Plethysmometer Model LE-7500 Plan lab S.L Italy), rotary evaporator Model-Heidolph HB digital, Germany, Vernier caliper, Nuclear magnetic resonance-Model EOL ECX 400 NMR, BOD Incubator-Model BI-81, Korea, UV-lamp.Autoclave-Model ST001045, Daihan scientific Co., Ltd. Korea, Laminar Flow Cabinet-Rashanica Pvt.Ltd, Hot plat model Hp-1D.

### 2.3 Test Animals Used

For the purpose of this research, albino mice of both sexes, with a weight range of 25-30 grams, were procured from the NIH (National Institute of Health), Islamabad, Pakistan. The subjects were housed in sanitary enclosures inside a controlled laboratory environment, characterized by a temperature range of (25± 2oC), humidity levels of (50±5oC), and a light/dark cycle lasting 12 hours. They were given a typical mouse foods and free availability of water *ad* libitum. Every aspect of animal research was conducted under strict human control. The experimental methodology was examined and has been authorized by the Research Ethical Committee of Department of Pharmacy, Abdul Wali Khan University, Mardan, Pakistan. The mice were only used once for the purpose of the research and then sacrificed according to protocol (Zeb et al., 2016).

### 2.4 Synthesis

A series of isoxazole derivatives were previously synthesized and characterization has been done (Alam et al., 2023).

### 2.5 5-LOX Inhibitory Assay

The synthesized Isoxazole derivatives will investigated for 5-LOX Inhibition studies as per established protocols of Wisastra and his co researchers (Wisastra et al., 2013). In this experiment, residual enzyme potential will be utilized to measure the degree of enzyme inhibition after 10 to 15 minutes of inhibitor’s incubation at 25°C. The reaction involving the conversion of linoleic acid, a substrate of lipoxygenase, into hydroperoxy-octadecadienoate (HPOD) will be employed as a means of quantifying the said reaction. The variation in the rate of absorption will be determined by employing a UV-visible spectrophotometer set at a wavelength of 234 nm. A 50 mM Tris buffer containing 2 mM EDTA and 2 mM CaCl_2_ was used as the assay buffer in this experiment. The buffer has a pH of 7.5. Following this, a buffer solution was employed to dilute the 5-lipoxygenase (5-LOX) enzyme at a concentration of 20,000 units per millilitre (U/mL) at a ratio of 1:4000. The 100 mM inhibitor was diluted using the test buffer following the administration of dimethyl sulfoxide (DMSO) for its dissolution. The content of linoleic acid was reduced to 20 mg/ml through dilution using ethyl alcohol. To accomplish this, 1 mL of the enzyme solution (1: 4000) will be combined with 100 µL of 2 mM adenosine triphosphate (ATP), 100 µL of the inhibitor (1 mM), and 790 mL of Tris buffer while incubated for 10 min. Following approximately 10 seconds of the substrate and the enzyme mixing together, 10 µL of a 20 mM substrate solution will be added to the mixture. At that point, the changing rate of the substrate will be determined. The response rate will be utilized as a positive control without blockers. In this experiment, montelukast will be utilized as the reference medicine.

### 2.6 DPPH Activity

The previously outlined methodology will be employed to evaluate the compounds’ antioxidant ability, as determined by the processes of the 1,1-diphenyl, 2-picrylhydrazyl (DPPH) free radical (Braca A et al., 2001). The experimental specimens will be subjected to dilution using different quantities, spanning from 62.5 to 1000 µg/ml, and afterwards introduced into a methanolic solution of DPPH at a concentration of 0.004%. At 517 nm, the absorbance will be determined by a UV spectrophotometer following a 30-minute duration. The scavenging capability of ascorbic acid will be quantified by employing the equation:

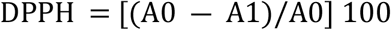

Whereas:

A0: Absorbance value of the control

A1: Absorbance value of the investigated compound concentration.

Every test was carried out three times, and the median inhibitory dosages (IC_50_) were determined. The drawing of inhibition graphs was performed by GraphPad Prism software-GraphPAD, San Diego, California, USA.

### 2.7 Statistical Analysis

The results of every investigation were presented as the mean SEM after being performed three times. A two-way ANOVA followed by Bonferroni posttest has been utilized to evaluate the positive control and investigated groups. *P* ˂0.05 has been used to predict the statistical significance.

## 3 RESULTS AND DISCUSSION

### 3.1 5-LOX Inhibitory Assay

The *in vitro* result findings of 5-LOX inhibitory potential of the designed tested derivatives and their IC_50_ values are computed in table 3.1. The *in vitro* 5-LOX investigations of the **C1** exhibited significant percent inhibition of 74.94±1.07, 71.39±0.60, 67.58±0.56, 62.29±1.43 and 56.37±0.58 at different concentration of 1000-62.5 µg/ml and the IC_50_ value determined for **C1** was 74.09 µM. Significant percent inhibition of 75.63±1.87, 71.12±0.54, 68.79±1.08, 63.79±1.88 and 58.20±0.47 were observed for **C2** at various concentrations with IC_50_ of 47.59 µM. The percent inhibition for **C3** were 89.93±1.73, 85.94±0.91, 81.90±1.32, 77.51±0.59 and 74.80±1.41 at various concentrations with IC_50_ of 8.47 µM. The percent inhibition of **C4** showed significant results having IC_50_ of 103.59 µM. The investigated compounds **C5** also exhibited good 5-LOX inhibitory effect. The non-significant percent inhibition for **C5** were demonstrated as 91.30±1.42, 87.78±0.45, 84.44±0.86, 79.72±1.89 and 75.29±1.64 at various concentration (1000-62.5 µg/ml). The IC_50_ demonstrated for **C5** was 10.48. Among the 10 synthesized compounds, the potential 5-LOX inhibitory effect was reported for **C6**. The non-significant percent inhibition observed at concentration 1000 µg/ml and 500 µg/ml for **C6** were 94.58±1.12 and 91.40±0.20 respectively, whereas the significant percent inhibition values reported at concentration 250, 125 and 62.5 µg/ml were 88.85±1.26, 85.08±0.47 and 81.90±0.96 respectively. The IC_50_ value found for **C6** was 3.67 µM. Similarly the assayed compound **C7** also exhibited non-significant percent inhibition of 94.80±0.90 at 1000 µg/ml and 90.94±1.13 at 500 µg/ml whereas significant percent inhibition was reported as 86.72±1.01, 81.84±0.30 and 78.80±1.50 at 250, 125 and 62.5 µg/ml, respectively. The percent inhibition for **C8** were established significant as 86.70±1.20 at 1000 µg/ml and 82.31±1.20 at 500 µg/ml whereas found non-significant as 78.30±2.67 at 250 µg/ml, 75.30±1.67 at 125 µg/ml and 71.03±1.79 at 62.5 µg/ml. The IC_50_ calculations noted for **C7** and **C8** were 10.51 and 9.80 µM, respectively. The remaining investigated compounds i.e. Compounds **C9** and **C10** also exhibited good activity against 5-LOX enzyme. The percent inhibition reported for **C9** were 81.42±0.43, 78.56±1.06, 74.90±2.45, 70.40±0.82, 67.33±1.66 and for **C10** were 73.31±0.35, 70.34±0.90, 66.78±0.34, 61.23±0.65, 55.34±1.34 respectively. The IC_50_ values determined for **C9** and **C10** were 11.25 and 67.06 µM, respectively. The percent inhibitions for montelukast as a standard drug was calculated as 94.08±1.04, 87.45±0.90, 81.58±1.63, 76.40±1.20 and 71.80±0.90 at 1000-62.5 µg/ml respectively. The IC_50_ values for montelukast was noted as 21.84 µM.

**Table 3.1.**
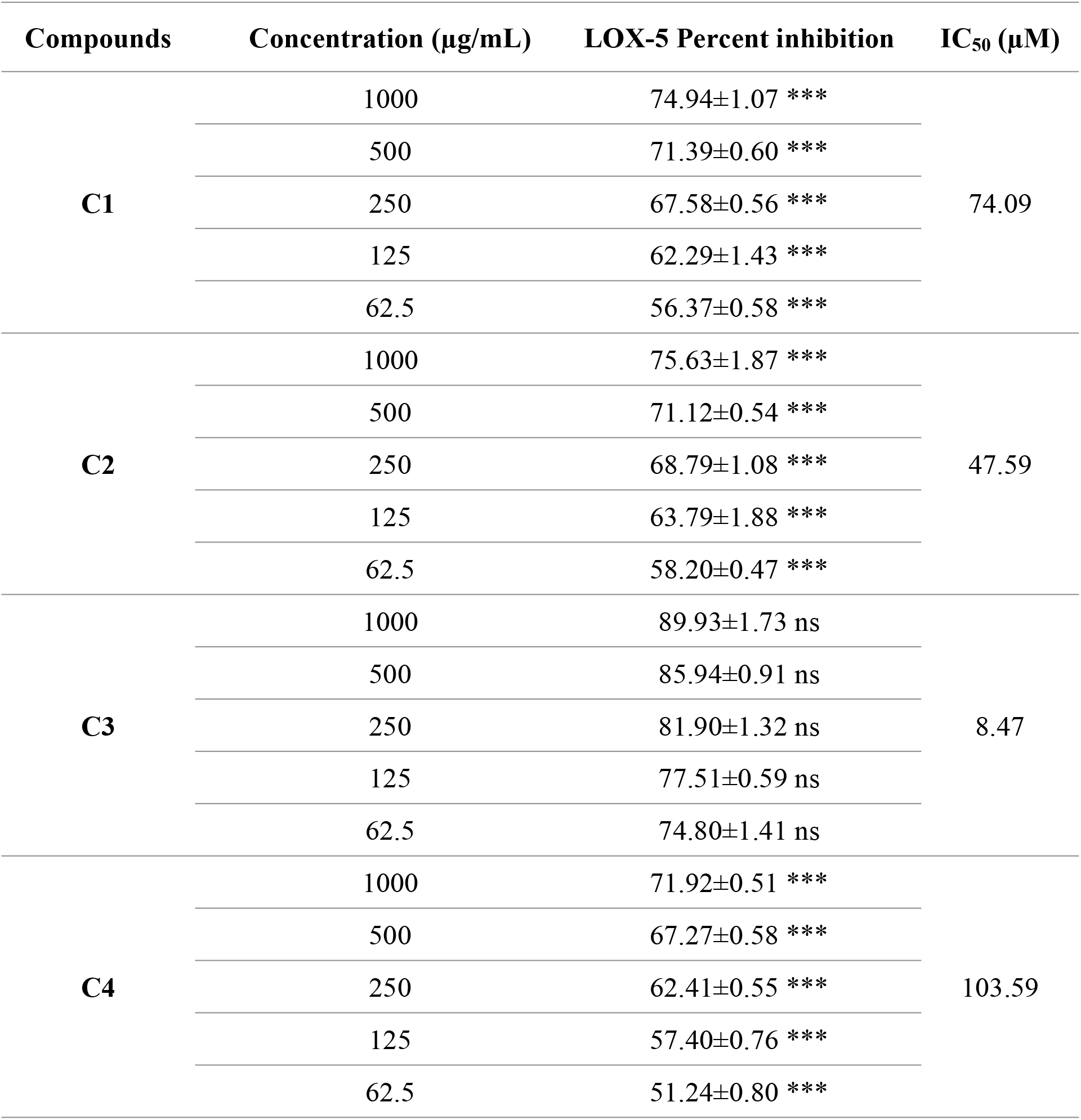
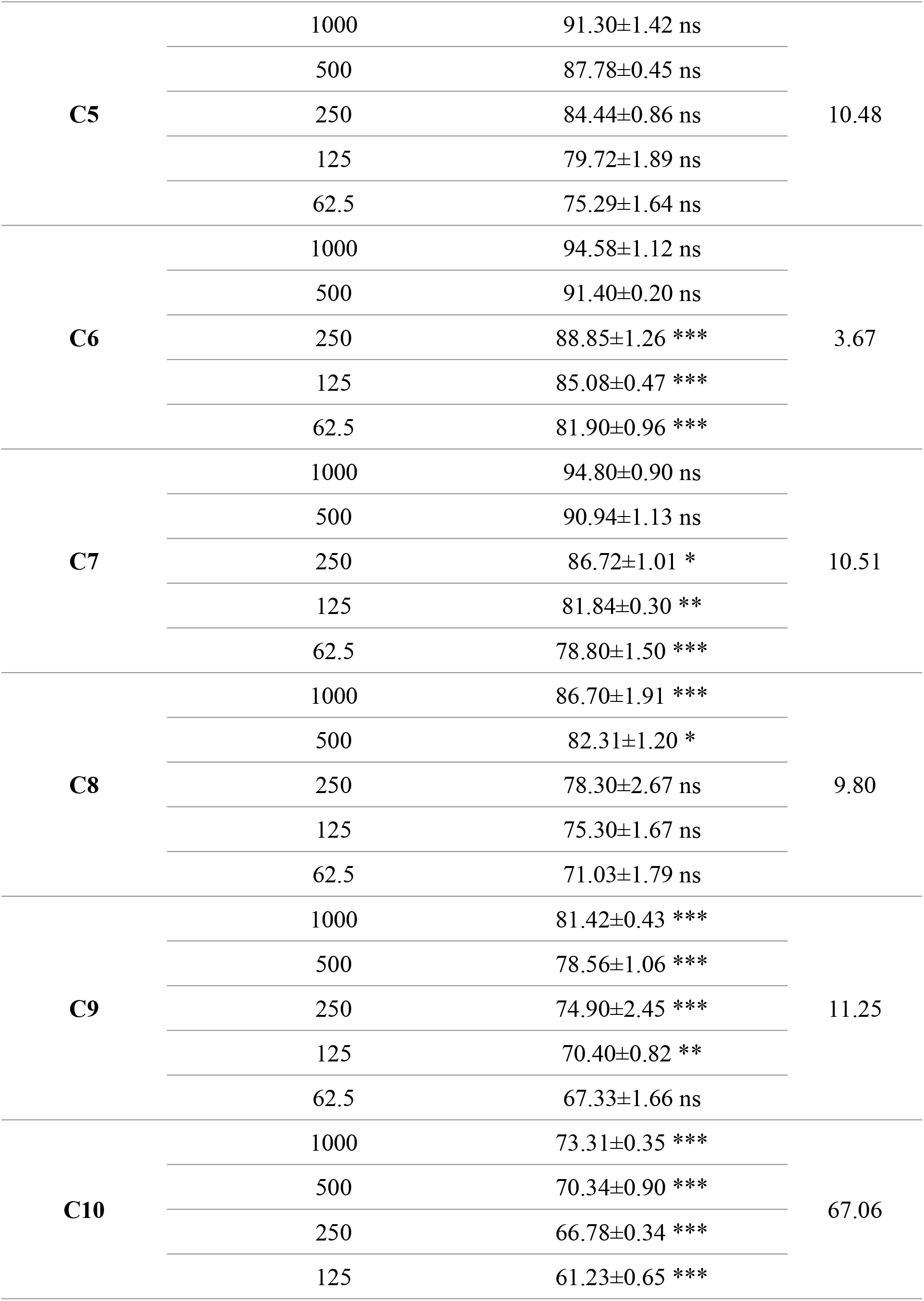
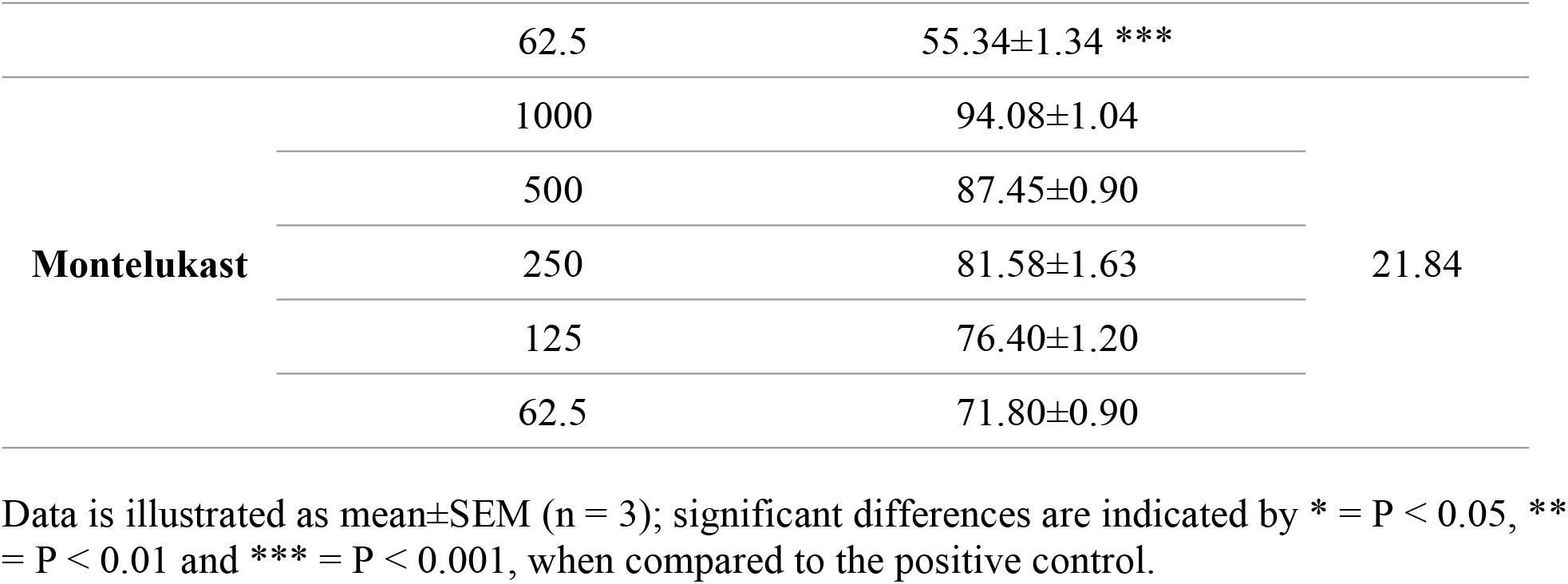
*In Vitro* 5-LOX Anti-inflammatory Activity of Synthesized Isoxazole Compounds against Montelukast as standard.

Lipoxygenases (LOXs) are non-heme iron-containing dioxygenases that generate hydroperoxy metabolites (HPETEs) by incorporating molecular oxygen into the substrate. There are six functioning LOX isoforms in humans (Srivastava et al., 2016). The 5-lipoxygenase (5-LOX) is an isozyme of LOX that initiates the second crucial metabolic process that produces eicosanoids. They are quite common in mammals, fungi, and plants. Most myeloid cells that are associated with inflammatory and immunological responses, which include polymorphonuclear leukocytes (eosinophils and neutrophils) and mononuclear cells (lymphocytes, monocytes, and macrophages), express the 5-LOX protein (Kaur and Silakari, 2017, Nguyen et al., 2020).

Leukotriene B4 is the final byproduct of the 5-LOX pathway and is a mediator of a number of inflammatory and allergic disorders, including atherosclerosis, cancer, and cardiovascular conditions. However, suppressing 5-LOX may aid to lessen the chance of cardiovascular and gastrointestinal problems triggered by COX-1/2 inhibitors, respectively, by lowering leukotriene levels. Notable is the possibility that COX/5-LOX co-inhibition may lessen adverse effects on the cardiovascular and gastrointestinal systems while maintaining the main effect of COX-1/2 inhibitors (Koukoulitsa et al., 2007, Martel-Pelletier et al., 2003).

Rakesh et al., has reported both the COX/5-LOX *in vitro* inhibitory effect of 3,5-Disubstituted Isoxazole compounds. They found that the preference for the target sites is largely determined by the presence of the pharmacophores (3-methylthienyl) and (3,4,5-trimethoxypheyl) in the side chains of the isoxazole. The inclusion of aryl moiety with strong electron-donor substituents and moderately electron-rich hetero aryl substituents (indole) may be the cause of compounds considerable inhibitory activity (Rakesh et al., 2016).

As previously researchers have reported that both *in vitro* inhibition of COX/5-LOX has shown excellent anti-inflammatory effects. Hence, in this study the *in vitro* 5-LOX enzyme inhibition potential of each tested compound was evaluated. The effectiveness of the tested analogues has been calculated as IC_50_ in µM that is the amount of the compound that results in 50% blocking of the enzyme. The result outcomes of our research established that the ten designed investigated isoxazole derivatives exhibited excellent 5-LOX blocking effect. The result findings of 5-LOX inhibition assays are represented in table 3.1. The 5-LOX enzymes inhibition analysis has been conducted for the tested derivatives at various concentration strengths. All the compounds were found potent and showed significant percent inhibition of 5-LOX pathway in dose dependent manner. Among the tested compounds, **C6** was found most potent which showed significant 5-LOX percent inhibition assay and also reported the minimum IC_50_ value comparable to the reference drug. The *in vitro* 5-Lox enzymes inhibition assays of **C5** and **C3** also showed non-significant percent inhibition and good potency next to **C6**. The IC_50_ values of **C5** and **C3** were also found effective. The remaining compounds also showed significant percent inhibition against 5-LOX enzyme inhibition assay.

### 3.2 DPPH Activity

The *in vitro* antioxidant effect of the synthesized derivatives has been evaluated against DPPH enzymes activity. The result findings are represented in table 3.10 and showed that almost all compounds has potent antioxidant effect against DPPH free radical scavenging activity. The most potent compound which showed excellent free radical scavenging effect was **C3** with different percent inhibitions of 92.51±0.62, 87.65±0.70, 82.25±0.55, 79.37±0.69 and 75.72±0.51 having IC_50_ value of 10.96 µM. The next most potent antioxidant activity was reported for **C5** which non-significantly showed free radical scavenging effect. The DPPH percent inhibition reported for **C5** was 92.63±0.64, 88.45±0.55, 83.53±0.41, 79.42±0.46 and 76.10±0.64 at different ranges of (1000, 500, 250, 125 and 62.5 µg/mL) concentrations, respectively. The IC_50_ value observed for **C5** was 13.12 µM. Compound **C6** also showed potent dose dependent antioxidant effect with IC_50_ value of 18.87 µM having percent inhibition of 91.63±0.55, 88.45±0.49, 83.53±0.45, 78.42±0.66 and 73.72±0.64 at concentration 1000-62.5 µg/mL respectively. The percent inhibition of **C1** and **C2** also expressed potent antioxidant effect against DPPH activity. At highest concentration (1000 µg/mL), the percent inhibition for **C1** and **C2** was 82.36±0.57 and 86.91±1.30 respectively. The reported IC_50_ value for **C1** and **C2** was 48.32 and 37.57µM respectively. Compound **C4** also showed significant free radicle scavenging effect against DPPH activity. The percent inhibition established for **C4** at highest concentration (1000 µg/mL) was 87.65±1.32 having IC_50_ value of 32.28 µM. The percent inhibition decreased to 65.03±0.48 at lowest concentration (62.5 µg/mL). The present investigation of DPPH activity for **C7, C8, C9** and **C10** also showed that a significant potent free radical scavenging effect was observed. The highest percent inhibition for **C7, C8, C9** and **C10** were 88.63 ± 1.57, 83.53±0.20, 81.85±0.18 and 88.88±0.89 respectively at highest concentration (1000 µg/mL) assay which decreased to 66.78 ± 0.72, 61.35±0.18, 59.12±0.34 and 59.82±0.95 respectively at lowest concentration (62.5 µg/mL). The IC_50_ values reported for **C7, C8, C9** and **C10** were 31.04, 54.71, 68.81 and 84.89 µM. Ascorbic acid, being used as a standard drug against DPPH antioxidant assay, demonstrated percent inhibition of 92.51±0.69, 87.65±0.42, 82.25±0.72, 79.37±0.71 and 77.72±0.59 at different concentration ranges of 1000-62.5 µg/mL respectively. The observed IC_50_ value for ascorbic acid was 17.90 µM.

The antioxidant potential of organic compounds must be evaluated as they are used in food, medicine, and cosmetics (Halliwell, 1996). Reactive species are produced in living systems by a variety of metabolic processes and stressful circumstances. They are primarily reactive oxygen species (ROS). Raised levels of ROS may alter biomolecules’ activities and impair their structural integrity, which can cause cellular malfunction and even premature death of cells. An rise in ROS over time can lead to oxidative stress at the systemic level, which manifests as a number of health issues including cancer, inflammation, age-related disorders, and heart problems (Grune et al., 2001, Noguchi and Niki, 2000). The most common and simple colorimetric technique for assessing the antioxidant capabilities of pure molecules is the DPPH assay, which is frequently used to determine how well a certain antioxidant molecule scavenges free radicals (Cheng et al., 2006). Researchers have reported antioxidant potential of isoxazole derivatives. Bhatia et al., has reported the antioxidant effect of Indole-Functionalized Isoxazoles derivatives. They found that in the DPPH test, the created compounds showed variable capacity to scavenge free radicals. The isoxazole ring of this molecule is considered as potent antioxidant moiety. The compounds’ capability to inhibit free radicals was significantly influenced by the substituted sequence at the phenyl ring connected to the isoxazole group (Bhatia et al., 2022). Yatoo and his co researchers reported the antioxidant potential of diosgenin based isoxazole derivatives. They found that the synthesized compounds exhibited highest antioxidant and anticancer effect (Yatoo and Banday, 2023).

In this *in vitro* research, the antioxidant potential of synthesized tested derivatives has been investigated in DPPH assay. The result findings of our antioxidant assay are represented in table 3.2. Our result outcomes established that almost all the tested compounds showed potent antioxidant effect against DPPH Activity. Various concentration ranges of the tested compounds were assayed against DPPH and their IC_50_ values were estimated as well. We concluded that amongst the investigated designed molecules the **C3** was found best potent and showed significant dose dependent antioxidant activity against DPPH screening. The IC_50_ value reported for **C3** was found good as compared to standard drug. Moreover, **C5** and **C6** also showed excellent free radical scavenging effect against DPPH assay. The remaining compounds also showed significant antioxidant effect. Based on the preliminary *in vitro* assays, the highest potent derivatives has been furtherly assayed for *in vivo* as well as histopathological screenings.

**Table 3.2.** *In Vitro* DPPH enzyme inhibition investigation of synthesized Isoxazole derivatives, using ascorbic acid as reference drug.

| Compounds | Mol.wt | Concentration (µg/mL) | DPPH Percent inhibition | IC<sub>50</sub> µM |
| --- | --- | --- | --- | --- |
| **C1** | 323.27 | 1000 | 82.36±0.57*** | 48.32 |
|  |  | 500 | 77.85±2.24*** |  |
|  |  | 250 | 72.08±0.47*** |  |
|  |  | 125 | 67.90±0.96*** |  |
|  |  | 62.5 | 62.28±0.57*** |  |
| **C2** | 353.3 | 1000 | 86.91±1.30*** | 37.57 |
|  |  | 500 | 81.26±1.27*** |  |
|  |  | 250 | 76.00±0.30*** |  |
|  |  | 125 | 69.54±0.50*** |  |
|  |  | 62.5 | 66.76±0.58*** |  |
| **C3** | 431.36 | 1000 | 92.51±0.62 ns | 10.96 |
|  |  | 500 | 87.65±0.70 ns |  |
|  |  | 250 | 82.25±0.55 ns |  |
|  |  | 125 | 79.37±0.69 ns |  |
|  |  | 62.5 | 75.72±0.51 ns |  |
| **C4** | 491.46 | 1000 | 87.65±1.32*** | 32.28 |
|  |  | 500 | 84.42±0.43* |  |
|  |  | 250 | 78.25±1.40** |  |
|  |  | 125 | 72.22±1.28*** |  |
|  |  | 62.5 | 65.03±0.48*** |  |
| **C5** | 365.26<br>(5.62) | 1000 | 92.63±0.64 ns | 13.12 |
|  |  | 500 | 88.45±0.55 ns |  |
|  |  | 250 | 83.53±0.41 ns |  |
|  |  | 125 | 79.42±0.46 ns |  |
|  |  | 62.5 | 76.10±0.64 ns |  |
| **C6** | 405.33<br>(7.165) | 1000 | 91.63±0.55 ns | 18.87 |
|  |  | 500 | 88.45±0.49 ns |  |
|  |  | 250 | 83.53±0.45 ns |  |
|  |  | 125 | 78.42±0.66 ns |  |
|  |  | 62.5 | 73.72±0.64** |  |
| **C7** | 443.3 | 1000 | 88.63 ± 1.57** | 31.04 |
|  |  | 500 | 84.17 ± 0.15** |  |
|  |  | 250 | 78.10 ± 0.14*** |  |
|  |  | 125 | 72.89 ± 0.17*** |  |
|  |  | 62.5 | 66.78 ± 0.72*** |  |
| **C8** | 389.33 | 1000 | 83.53±0.20*** | 54.71 |
|  |  | 500 | 78.62±0.17*** |  |
|  |  | 250 | 73.42±0.11*** |  |
|  |  | 125 | 66.20±0.15*** |  |
|  |  | 62.5 | 61.35±0.18*** |  |
| **C9** | 386.33 | 1000 | 81.85±0.18*** | 68.81 |
|  |  | 500 | 76.59±0.30*** |  |
|  |  | 250 | 69.75±0.14*** |  |
|  |  | 125 | 64.47±0.49*** |  |
|  |  | 62.5 | 59.12±0.34*** |  |
| **C10** | 387.32 | 1000 | 88.88±0.89** | 84.78 |
|  |  | 500 | 83.54±0.60*** |  |
|  |  | 250 | 75.01±0.97*** |  |
|  |  | 125 | 67.68±0.22*** |  |
|  |  | 62.5 | 59.82±0.95*** |  |
| **Ascorbic acid** | 176.12 | 1000 | 92.51±0.69 | 17.90 |
|  |  | 500 | 87.65±0.42 |  |
|  |  | 250 | 82.25±0.72 |  |
|  |  | 125 | 79.37±0.71 |  |
|  |  | 62.5 | 77.72±0.59 |  |

Data is illustrated as mean±SEM (n = 3); significant differences are indicated by \* = P < 0.05, \*\* = P < 0.01 and \*\*\* = P < 0.001, when compared to the positive control.

## 4 Conclusion

The 5-LOX enzymes inhibition analysis has been conducted for the tested derivatives at various concentration strengths. All the compounds were found potent and showed significant percent inhibition of 5-LOX pathway in dose dependent manner. Among the tested compounds, **C6** was found most potent which showed significant 5-LOX percent inhibition assay and also reported the minimum IC_50_ value comparable to the reference drug. The *in vitro* 5-Lox enzymes inhibition assays of **C5** and **C3** also showed non-significant percent inhibition and good potency next to **C6**. Our result outcomes established that almost all the tested compounds showed potent antioxidant effect against DPPH Activity. Moreover, **C5** and **C6** also showed excellent free radical scavenging effect against DPPH assay. The remaining compounds also showed significant antioxidant effect. Based on the preliminary *in vitro* assays, the highest potent derivatives has been furtherly assayed for *in vivo* as well as histopathological screenings.

## References

Alam, W., Khan, H., Jan, M. S., Rashid, U., Abusharha, A. & Daglia, M. 2023. Synthesis, in-vitro inhibition of cyclooxygenases and in silico studies of new isoxazole derivatives. Frontiers in Chemistry, 11.

Bhatia, R., Vyas, A., El-Bahy, S. M., Hessien, M. M., Mersal, G. A., Ibrahim, M. M., Dogra, R. & Kumar, B. 2022. Rationale Design, Synthesis, Pharmacological and In-silico Investigation of Indole-Functionalized Isoxazoles as Anti-inflammatory Agents. ChemistrySelect, 7, e202200800.

Biringer, R. G. 2020. The enzymology of human eicosanoid pathways: The lipoxygenase branches. Molecular Biology Reports, 47, 7189–7207.

Braca A, Tommasi ND, Bari LD, Pizza C, M, P. & and Morelli, I. 2001. Antioxidant principles from Bauhinia terapotensis. Journal of Natural Products, 64, 892–895.

Cheng, Z., Moore, J. & Yu, L. 2006. High-throughput relative DPPH radical scavenging capacity assay. Journal of Agricultural and Food Chemistry, 54, 7429–7436.

FeißT, C., Pergola, C., Rakonjac, M., Rossi, A., Koeberle, A., Dodt, G., Hoffmann, M., Hoernig, C., Fischer, L. & Steinhilber, D. 2009. Hyperforin is a novel type of 5-lipoxygenase inhibitor with high efficacy in vivo. Cellular and molecular life sciences, 66, 2759–2771.

Fischer, L., Hornig, M., Pergola, C., Meindl, N., Franke, L., Tanrikulu, Y., Dodt, G., Schneider, G.,Steinhilber, D. & Werz, O. 2007. The molecular mechanism of the inhibition by licofelone of the biosynthesis of 5-lipoxygenase products. British journal of pharmacology, 152, 471–480.

Gaschler, M. M. & Stockwell, B. R. 2017. Lipid peroxidation in cell death. Biochemical and biophysical research communications, 482, 419–425.

Grune, T., Shringarpure, R., Sitte, N. & Davies, K. 2001. Age-related changes in protein oxidation and proteolysis in mammalian cells. The Journals of Gerontology Series A: Biological Sciences and Medical Sciences, 56, B459–B467.

Halliwell, B. 1996. Antioxidants in human health and disease. Annual Review of Nutrition, 16, 33–50.

Hofmann, B. & Steinhilber, D. 2013. 5-Lipoxygenase inhibitors: a review of recent patents (2010–2012). Expert opinion on therapeutic patents, 23, 895–909.

Kaur, G. & Silakari, O. 2017. Multiple target-centric strategy to tame inflammation. Future Medicinal Chemistry, 9, 1361–1376.

Koukoulitsa, C., Hadjipavlou–Litina, D., Geromichalos, G. D. & Skaltsa, H. 2007. Inhibitory effect on soybean lipoxygenase and docking studies of some secondary metabolites, isolated from Origanum vulgare L. ssp. hirtum. Journal of Enzyme Inhibition and Medicinal Chemistry, 22, 99–104.

Lenis, J. M., Gillman, J. D., Lee, J. D., Shannon, J. G. & Bilyeu, K. D. 2010. Soybean seed lipoxygenase genes: molecular characterization and development of molecular marker assays. Theoretical and applied genetics, 120, 1139–1149.

Martel-Pelletier, J., Lajeunesse, D., Reboul, P. & Pelletier, J.-P. 2003. Therapeutic role of dual inhibitors of 5-LOX and COX, selective and non-selective non-steroidal anti-inflammatory drugs. Annals of the Rheumatic Diseases, 62, 501–509.

Mcmillan, R. & Walker, E. 1992. Designing therapeutically effective 5-lipoxygenase inhibitors. Trends in pharmacological sciences, 13, 323–330.

Muthuraman, S., Sinha, S., Vasavi, C., Waidha, K. M., Basu, B., Munussami, P., Balamurali, M., Doble, M. & Kumar, R. S. 2019. Design, synthesis and identification of novel coumaperine derivatives for inhibition of human 5-LOX: Antioxidant, pseudoperoxidase and docking studies. Bioorganic & Medicinal Chemistry, 27, 604–619.

Nguyen, H. T., Vu, T.-Y., Chandi, V., Polimati, H. & Tatipamula, V. B. 2020. Dual COX and 5-LOX inhibition by clerodane diterpenes from seeds of Polyalthia longifolia (Sonn.) Thwaites. Scientific Reports, 10, 1–10.

Noguchi, N. & Niki, E. 2000. Phenolic antioxidants:: A rationale for design and evaluation of novel antioxidant drug for atherosclerosis. Free Radical Biology and Medicine, 28, 1538–1546.

Prasad, N. S., Raghavendra, R., Lokesh, B. & Naidu, K. A. 2004. Spice phenolics inhibit human PMNL 5-lipoxygenase. Prostaglandins, leukotrienes and essential fatty acids, 70, 521–528.

Rakesh, K. S., Jagadish, S., Balaji, K. S., Zameer, F., Swaroop, T. R., Mohan, C. D., Jayarama, S. & Rangappa, K. S. 2016. 3, 5-Disubstituted isoxazole derivatives: Potential inhibitors of inflammation and cancer. Inflammation, 39, 269–280.

Sinha, S., Sravanthi, T., Yuvaraj, S., Manju, S. & Doble, M. 2016. 2-Amino-4-aryl thiazole: a promising scaffold identified as a potent 5-LOX inhibitor. RSC advances, 6, 19271–19279.

Srivastava, P., Vyas, V. K., Variya, B., Patel, P., Qureshi, G. & Ghate, M. 2016. Synthesis, anti-inflammatory, analgesic, 5-lipoxygenase (5-LOX) inhibition activities, and molecular docking study of 7-substituted coumarin derivatives. Bioorganic Chemistry, 67, 130–138.

Tomy, M. J., Sharanya, C. S., Dileep, K. V., Prasanth, S., Sabu, A., Sadasivan, C. & Haridas, M. 2015. Derivatives form better lipoxygenase inhibitors than piperine: in vitro and in silico study. Chemical Biology & Drug Design, 85, 715–721.

Wisastra, R., Kok, P. A., Eleftheriadis, N., Baumgartner, M. P., Camacho, C. J., Haisma, H. J. & Dekker, F. J. 2013. Discovery of a novel activator of 5-lipoxygenase from an anacardic acid derived compound collection. Bioorganic & medicinal chemistry, 21, 7763–7778.

Yatoo, G. N. & Banday, J. A. 2023. Synthesis, antioxidant, antiproliferative activity, molecular docking and DFT studies of novel isoxazole derivatives of diosgenin, a steroidal sapogenin from Dioscorea deltoidea. Fitoterapia, 105621.

Zeb, A., Ahmad, S., Ullah, F., Ayaz, M. & Sadiq, A. 2016. Anti-nociceptive activity of ethnomedicinally important analgesic plant Isodon rugosus Wall. ex Benth: Mechanistic study and identifications of bioactive compounds. Frontiers in pharmacology, 7, 200.

